# Isolation and characterization of BpL1, a broad acting lytic bacteriophage against Brucella

**DOI:** 10.1101/165076

**Authors:** Vimlesh Gupta, Hari Mohan Saxena

## Abstract

We have isolated a new broad acting lytic brucellaphage (BpL1) from the sewage of a dairy farm. The phage lysed all the 12 *Brucella abortus* field isolates, *B. abortus* strain 99 and *Brucella melitensis* but did not lyse any of the heterologous species tested viz. *Staphylococcus aureus, Pasteurella multocida, Escherichia coli,* and *Salmonella species*. Streaking the plaques on *Brucella* lawn gave clear zones along the streak lines. The plaques were circular with a diameter of 0.5- 3.0 mm. At a concentration of 10^−4^ the phage count was 4.5 × 10^6^ plaques per ml. It was a tailed phage with icosahedral head (62.2 nm in diameter and 73.71 nm in length), and the head to tail length was 229.21 nm. The phage belonged to the order Caudovirales and family *Siphoviridae*. It was inactivated within one hour at 55°C and within 4 hours at −20°C. Treatment at pH 2 for 4 hours and at pH 4 for 12 hours inactivated it. It was inactivated after 4 hours exposure to sunlight, and within 4 minutes by UV light. Chloroform and Sodium Dodecyl Sulfate inactivated it within 15 minutes. Lysozyme inactivated it within 1 hour whereas RNase treatment did not affect its activity.

## Introduction

Brucellosis is a highly contagious and important zoonotic disease caused by different species of the genus Brucella. The Brucella organisms are pathogenic for a wide variety of animals such as swine, cattle, goat, sheep, and dogs and also for humans (Mathur *et al* 2007).In animals, brucellosis mainly affects reproduction and fertility, reduces the survival of newborns, and diminishes milk yield. Outbreak of brucellosis in animals is characterized by abortions during the last trimester of gestation. Brucellosis is endemic in India and is prevalent in all parts of the country. In Punjab, prevalence of brucellosis has been found to be 17.68% (Aulakh *et al* 2008) and is considered the major cause of abortion in animals. Currently, no effective and affordable treatment of Brucellosis in large animals is available. Antibiotic resistance has been detected in Brucella organisms also. The widely used S-19 vaccine has not been successful in controlling Bucellosis since infection in some vaccinated animals has also been observed (Mohan *et al*., 2016). A bacteriophage is a virus that specifically infects and lyses its host bacterium. This unique characteristic can be exploited for therapy of infections due to antibiotic resistant bacteria. We report here the isolation of a new Brucellaphage from the habitat of Brucella infected animals i.e. sheds of infected cattle.

## Materials and methods

### Isolation, identification and characterization of *Brucella abortus*

*Brucella abortus* vaccine strain S19, *Brucella abortus* S99, *B. melitensis* Rev1 and *B. melitensis* were grown on Brucella selective agar (HiMedia) and incubated at 37°C aerobically. *B. abortus* from field samples were isolated on Brucella selective agar. Samples comprising of aborted foetal stomach contents, placenta and cotyledons and vaginal and uterine secretions from cattle and buffaloes with a history of abortions were collected and were subjected to isolation. *Brucella* isolates were identified on the basis of cultural, morphological and biochemical characteristics. DNA extraction was done as per the standard protocols of Sambrook and Russell (2001). Molecular characterization of the field isolates were done by polymerase chain reaction (PCR) as per the method of Romero *et al*. (1995).

### Isolation of Bacteriophage against *Brucella abortus*

Agar overlay technique was used to isolate bacteriophage against *Brucella abortus* (Adams 1959 and Chilamban 2004). A total of seven sewage samples were collected from a dairy farm at different times and processed for the isolation of phage. In brief, to the 50 ml double strength NZCYM broth (Life Technologies) 40 ml sewage supernatant and 10 ml of broth culture of *B. abortus* in exponential growth were added and incubated on rotary shaker for 10 days at 37°C. Out of this incubated sewage + bacteria cocktail, 10 ml of supernatant was taken every day and centrifuged at 8000g for 15 minutes to collect the supernatant which was passed through 0.22μm PVDF filter (Axiva) and the filtrate was aseptically collected and stored at 4°C till further use and was designated as Bacteria Free Filtrate (BFF). Equal quantities (100 μl) of BFF and overnight broth culture of *B. abortus* were mixed in 0.75% NZCYM agar (maintained at 45°C in a dry bath) and was spread evenly over 1.5% NZCYM agar + BSM agar. The soft agar was allowed to solidify and the plates were incubated at 37°C for 48-72 h to observe plaques.

### Elution of Brucellaphage

The plaques were picked using a straight wire loop and were streaked horizontally and then vertically on a hardened NZCYM + BSM plate overlaid with semisolid NZCYM agar containing the indicator strain. The plate was incubated at 37°C for 18h to observe plaques along the lines. SM buffer (2ml) was poured over the agar and the agar was disturbed with the wire loop to release the phages from the semisolid agar. This SM buffer was then collected and centrifuged at 5000g to remove the agar and then the supernatant was filtered through 0.22 μm filters to remove the bacteria and elute the phage in SM buffer.

### Heterologous species of bacteria used for testing the phage

A set of heterologous species of bacteria of veterinary importance viz. *Pasteurella multocida B: 2, E.coli, Staphylococcus, Streptococcus, Salmonella Dublin, Micrococcus* and *Pseudomonas spp* available in the Department of Veterinary Microbiology, GADVASU were used.

### Effect of varied temperatures on the brucellaphage

100μl of brucellaphage (10^6^pfu/ml) was subjected to temperatures of −20°C, 4°C, 37°C, 50°C, 70°C and 100°C for a period of 20 min. Any change in pfu was observed by adding 200μl of freshly grown *Brucella abortus S19* culture in cooled molten semisolid NZCYM and plated on BSM+NZCYM plates. These plates were incubated aerobically at 37°C for 48 to 72 h.

### Effect of light on the brucellaphage

100μl of brucellaphage (10^6^pfu/ml) was subjected to normal fluorescent tube light, sunlight and UV light for a period of 15 min to 90 min. Any change in pfu was observed by adding 200μl of freshly grown *Brucella abortus S19* culture in cooled molten semisolid NZCYM and plated on BSM+NZCYM plates. These plates were incubated aerobically at 37°C for 48 to 72 h.

### Effect of enzymes on the brucellaphage

100μl of brucellaphage (10^6^pfu/ml) and 100 μl of enzymes *viz*. proteinase K (20mg/ml), trypsin (250μg/ml), lysozyme (20mg/ml) and RNAse (10mg/ml) were incubated for 15 min. Any change in pfu was observed by adding 200μl of freshly grown *Brucella abortus S19* culture in cooled molten semisolid NZCYM and plated on BSM+NZCYM plates. These plates were incubated aerobically at 37°C for 48 to 72 h.

### Effect of SDS (10%), NSS, EDTA (0.01M) on the brucellaphage

100μl of brucellaphage (10^6^pfu/ml) was subjected to treatment with equal volume of SDS(10%), NSS and EDTA(0.01M) for a period of 15 min to 3h. Any change in pfu was observed by adding 200μl of freshly grown *Brucella abortus S19* culture in cooled molten semisolid NZCYM and plated on BSM+NZCYM plates. These plates were incubated aerobically at 37°C for 48 to 72 h.

### Effect of varied pH on the survivability of bacteriophage

100μl of brucellaphage (10^6^pfu/ml) was observed for the change in pfu count in different pH ranges of 3, 5, 7 and 9 for a period of 30 min exposure time and 60 min exposure time. Any change in pfu was observed by adding 200μl of freshly grown *Brucella* culture in cooled molten semisolid NZCYM and plated on BSM+NZCYM plates. These plates were incubated aerobically at 37°C for 48 to 72 h.

## Results and discussion

### Isolation of brucellaphage

Out of the total 36 sewage samples, brucellaphage could be isolated from 1 sample. Streaking the plaques on *Brucella* lawn gave clear zones along the streak lines (Fig 1).

**Fig 1.**
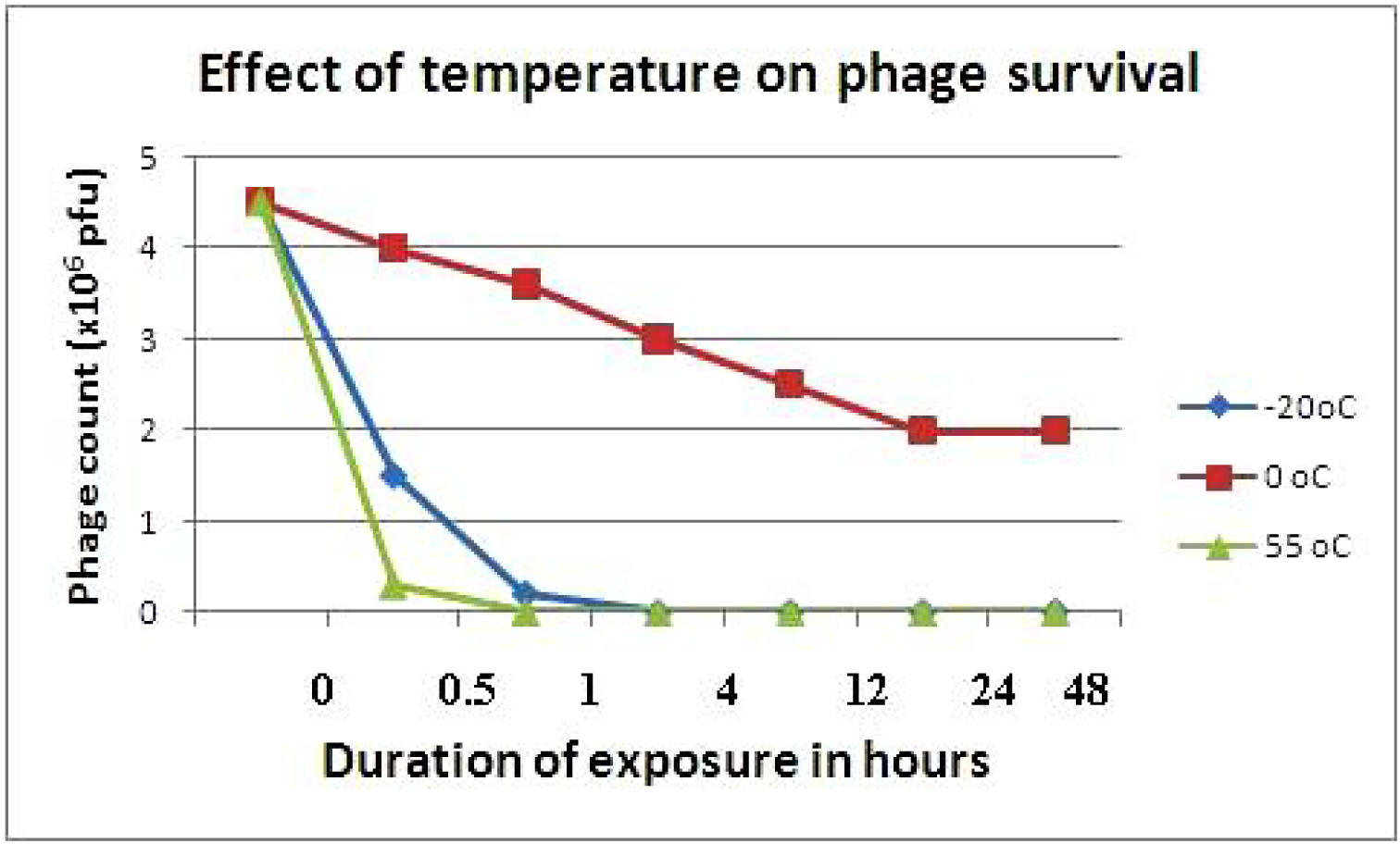
Effect of temperature on phage survival.

### Plaque forming unit (pfu/ml)

At a concentration of 10^−4^ the phage count was 4.5 × 10^6^ plaques per ml.

### Heterogenecity test for brucellaphage

The isolated brucellaphage lysed all the 12 *Brucella abortus* field isolates, *B. abortus* strain 99 and *Brucella melitensis* (procured from IVRI, Izatnagar) but did not lyse any of the heterologous species tested viz. *Staphylococcus aureus, Salmonella species, Escherichia coli,* and *Pasteurella multocida*. These observations were similar to those of Chachra *et al* (2012) Pandey *et al* (2013) and who had reported that their brucellaphage did not lyse any of the heterologous bacterial species tested *viz*. *Staphylococcus aureus, Streptococcus* species, *E. coli, P. multocida, Pseudomonas* species and *Salmonella* Dublin. Morris and Corbel (1973) have reported that the isolated Weybridge phage was lytic for smooth *B. abortus* biotype (1, 2, 3, 4, 5, 6, 7 and 9), *B. suis* biotype (1, 2, 3 and 4), and *B. neotomae* cultures but there was no lysis of *B. canis, B. melitensis* and *B. ovis* strains.

## Morphological characterization of brucellaphage

### Plaque morphology

The observed plaques were circular with a diameter of 0.5- 3.0 mm (Fig 2). Pandey *et al* (2013) had reported clear plaques of brucellaphage of variable size (0.1 to 3 mm). Morris *et al* (1973) observed clear plaques of phage A422 of 0.1 to 2.0 mm diameter whereas the plaques of S708 and M51 were of two types i.e. small, turbid plaques, 0.1 to 0.5 mm in diameter, and large, clear plaques, 0.5 to 2.5 mm in diameter.

**Fig 2.**
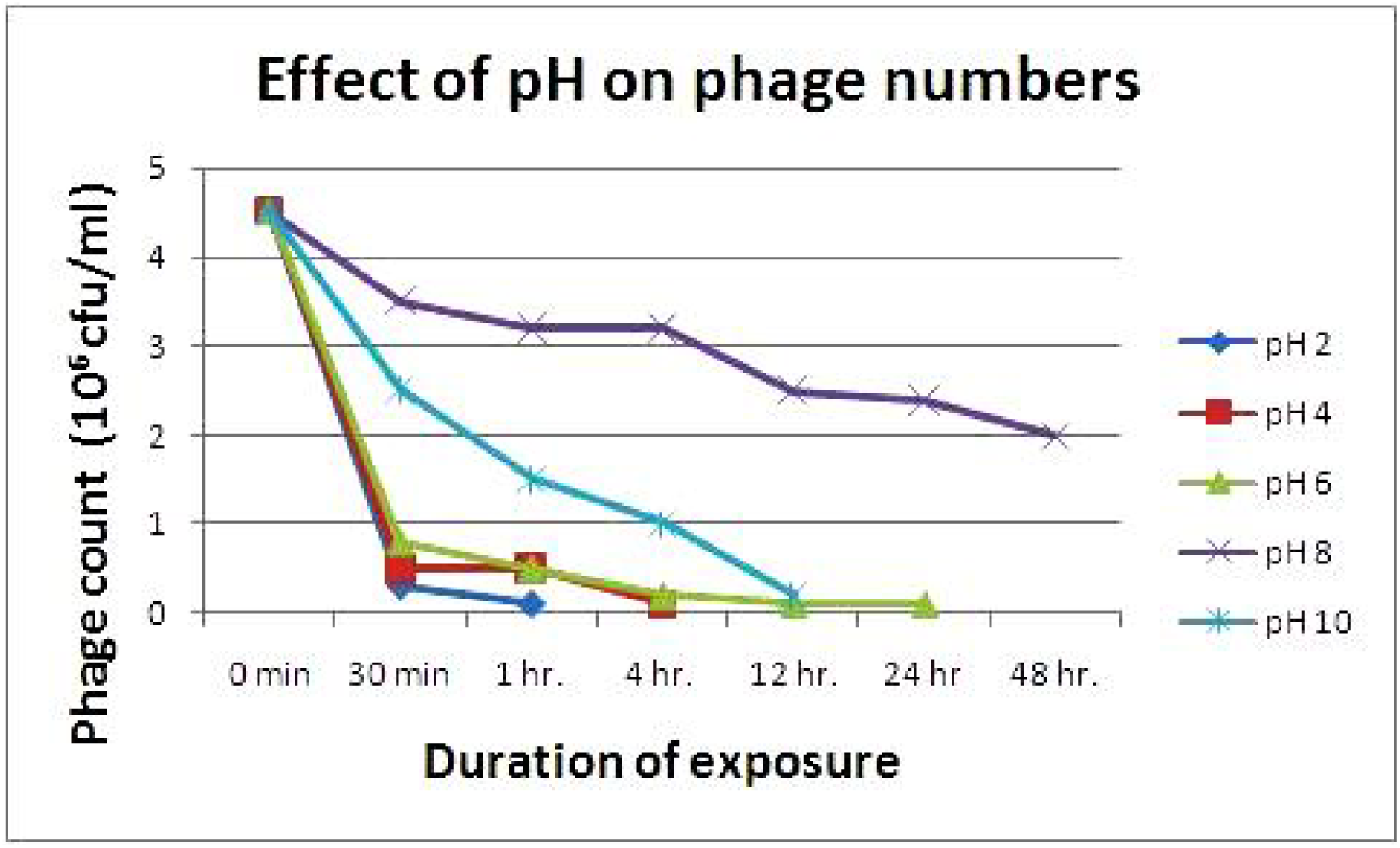
Effect of pH on phage survival.

### Phage morphology

The brucellaphage isolated in the present study was a tailed phage with icosahedral head (62.2 nm in diameter and 73.71 nm in length), and the head to tail length was 229.21 nm (Fig 3). According to Ackermann (2007) such types of phages belong to the order Caudovirales and family *Siphoviridae*. We propose to name it as “Brucellaphage Ludhiana 1 (BpL1)”.

**Fig 3:**
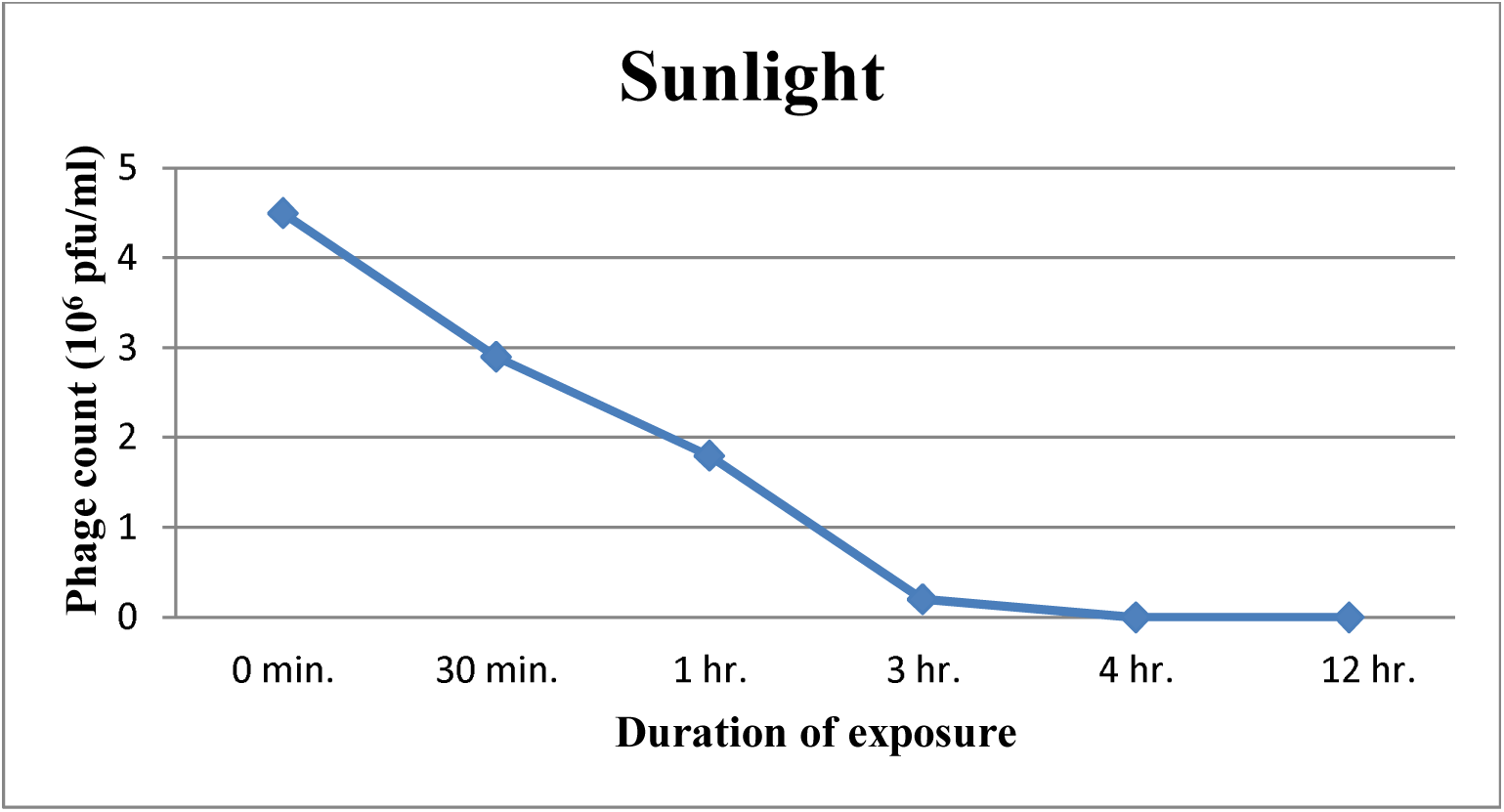
Effect of exposure to sunlight on brucellaphage survivability.

Cai-Zhong *et al* (2009) classified Tb (*Tbilisi*), as a member of the *Podoviridae* family with icosahedral capsids (57 ± 2 nm diameter) and short tails (32 ± 3 nm long). Chachra *et al* (2012) reported that electron microscopic studies of the brucellaphage revealed it to be an elementary body measuring approximately 65 nm with rounded head and a very short tail. Rigby *et al* (1989) reported that Nepean (Np) was morphologically identical to the other brucellaphages, with an icosahedral head (diameter 50-65 nm) and short tail (length 15-20 nm).

## Physicochemical characterization of the brucellaphage

### Effect of temperature

The effect of temperature on the survivability of brucellaphage was studied. The phage titre gradually decreased from 4.5 × 10^6^ to 2.0 × 10^6^ pfu/ml within 48 hours at 0°C. Temperature treatments of −20°C completely inactivated the phage within 4 hours and treatment at 55°C completely inactivated the phage within 1 hour (Table 1, fig 1).

**Table 1:**
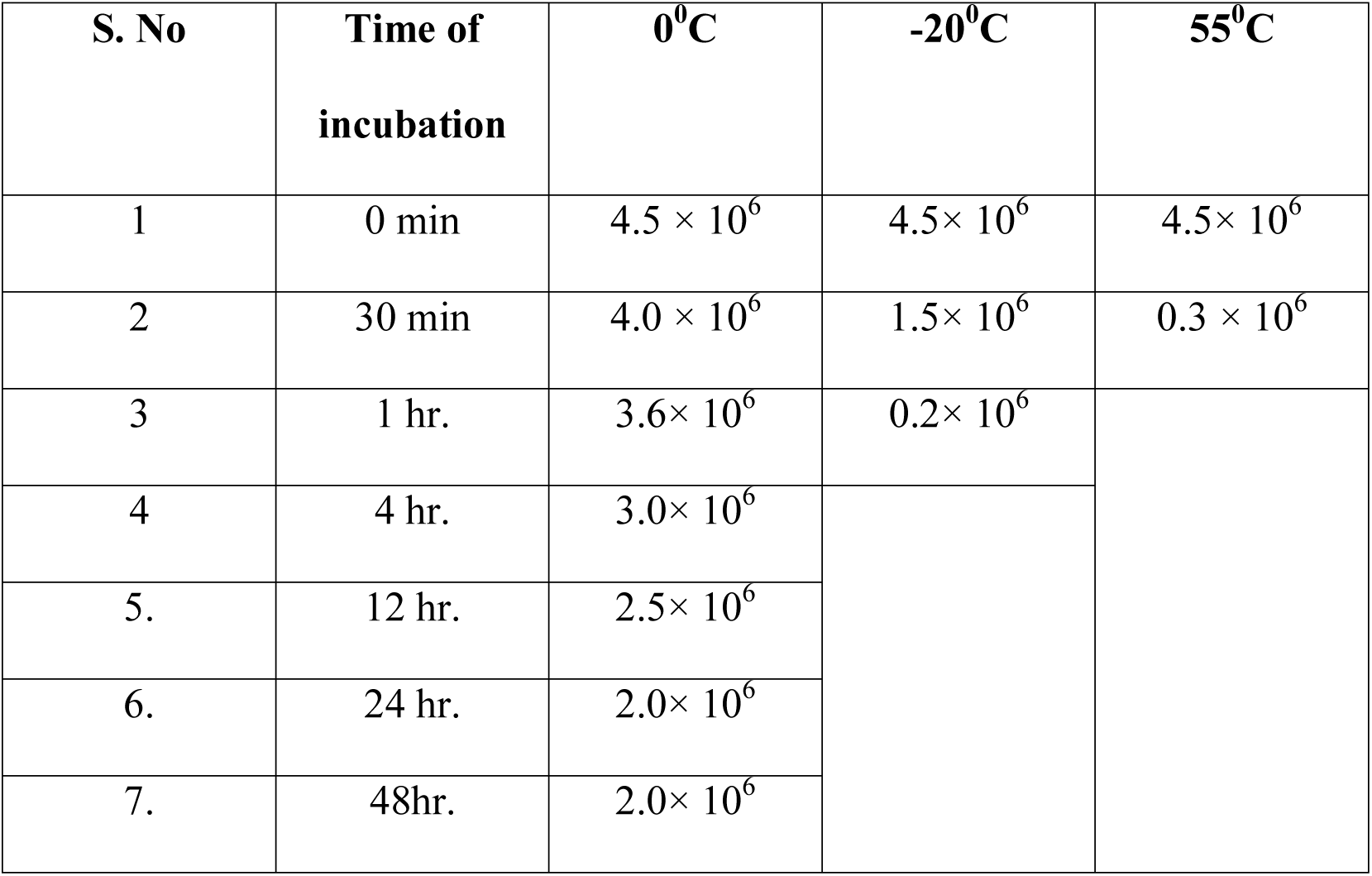
Effect of temperature on survivability of brucellaphage.

McDuff *et al* (1961) had reported that 18% brucellaphages were inactivated at 60°C and there was 100% inactivation at 70°C within 60 minutes. Pandey *et al* (2013) have also reported that at 40°C the brucellaphage titre gradually decreased from 1340 pfu/ml to 110 pfu/ml within 3 hours and treatment at 60°C completely inactivated the phage within 10 minutes.

### Effect of pH on brucellaphage

Various pH treatments (pH 2, 4, 6, 8, 10) were given to the brucellaphage to determine its survivability. The phage was inactivated at pH 2 within 4 hours, and at pH 4 within 12 hours. However the phage survived up to 24 hours at pH 6.The phage number gradually decreased from 4.5× 10^6^ to 2.0× 10^6^ within 48 hours at pH 8. It decreased to 0 within 24 hours at pH 10 (Table 2, Fig 2). These observations were similar to the observations of McDuff *et al* (1961) who reported that there was no loss in phage titre in broth at pH values of 6.2 to 8.1 whereas, at pH 3.1 there was complete (100%) inactivation, 56% at pH 4.1, 24% at pH 5.0, 35% at pH 9.0 and 42% inactivation at pH 9.9, respectively. Pandey *et al* (2013) reported that treatment at pH 2 completely inactivated the phage within 3 h, whereas the phage titre gradually decreased to zero within 24 h at pH 4. At pH 6, there was only 38.9% decrease in the phage titre after 48 h of treatment. The phage remained stable at pH 8 with 75.31% survivability after 48 h treatment. At pH 10, the phage titre gradually decreased to 0.31% within 48 h.

**Table 2:** Effect of pH on brucellaphage survivability.

| S. No | Time duration | pH 2 | pH 4 | pH 6 | pH 8 | pH 10 |
| --- | --- | --- | --- | --- | --- | --- |
| 1 | 0 min | $4.5 \times 10^6$ | $4.5 \times 10^6$ | $4.5 \times 10^6$ | $4.5 \times 10^6$ | $4.5 \times 10^6$ |
| 2 | 30 min | $0.3 \times 10^6$ | $0.5 \times 10^6$ | $0.8 \times 10^6$ | $3.5 \times 10^6$ | $2.5 \times 10^6$ |
| 3 | 1 hr. | $0.1 \times 10^6$ | $0.5 \times 10^6$ | $0.5 \times 10^6$ | $3.2 \times 10^6$ | $1.5 \times 10^6$ |
| 4 | 4 hr. | | $0.1 \times 10^6$ | $0.2 \times 10^6$ | $3.2 \times 10^6$ | $1.0 \times 10^6$ |
| 5. | 12 hr. | | | $0.1 \times 10^6$ | $2.5 \times 10^6$ | $0.2 \times 10^6$ |
| 6. | 24 hr. | | | $0.1 \times 10^6$ | $2.4 \times 10^6$ | |
| 7 | 48 hr. | | | | $2.0 \times 10^6$ | |

### Effect of sunlight

When the brucellaphage was exposed to sunlight, the phage titre decreased gradually from 4.5 × 10^6^ to 0.2 × 10^6^ within 3 hours and was completely inactivated after 4 hours (Table 3, fig 3). Pandey *et al* (2013) have also reported that direct sunlight gradually decreases the phage titre and within 3 hours brucellaphage is reduced by 93.99%.

**Table 3:**
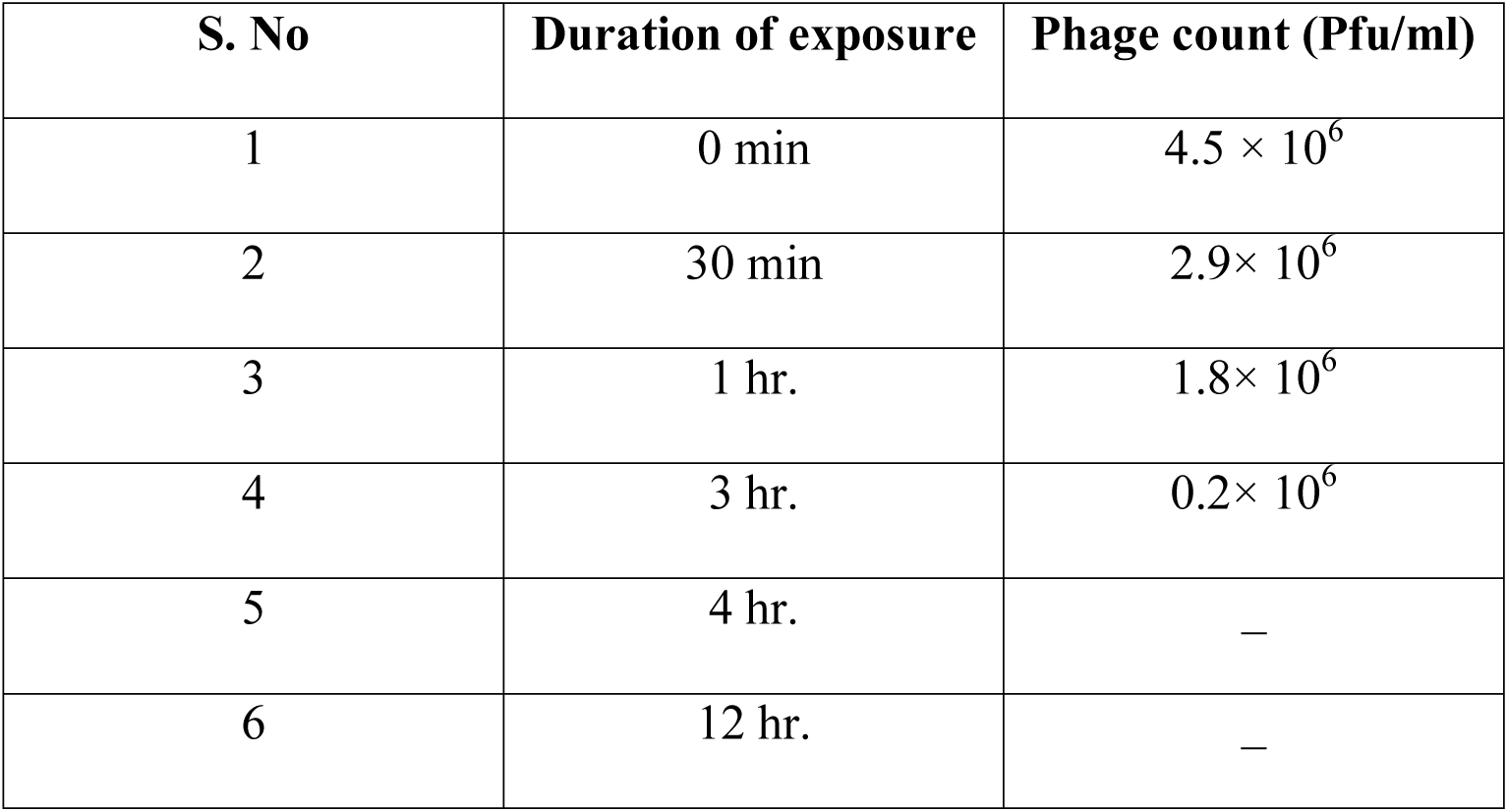
Effect of sunlight on brucellaphage survival.

### Effect of UV light

Exposure to UV light was found to have a drastic effect on phage survivability. The phage gets completely inactivated within 4 minutes (Table 20, fig 14). Pandey *et al* (2013) had also reported that exposure to UV rays inactivated the phage completely within 3 minutes.

## Chemical characterization of phage

### Effect of Chloroform (10%) on phage

The treatment of phage with 10% chloroform completely inactivated it within 15 minutes at 37°C (Table 5, fig 5). McDuff *et al* (1961) had reported that there was 99% inactivation of phage by 10% chloroform within 5 minutes. Pandey *et al* (2013) had also reported the complete inactivation within 5 minutes of exposure of phage to 10% chloroform.

**Table 5:**
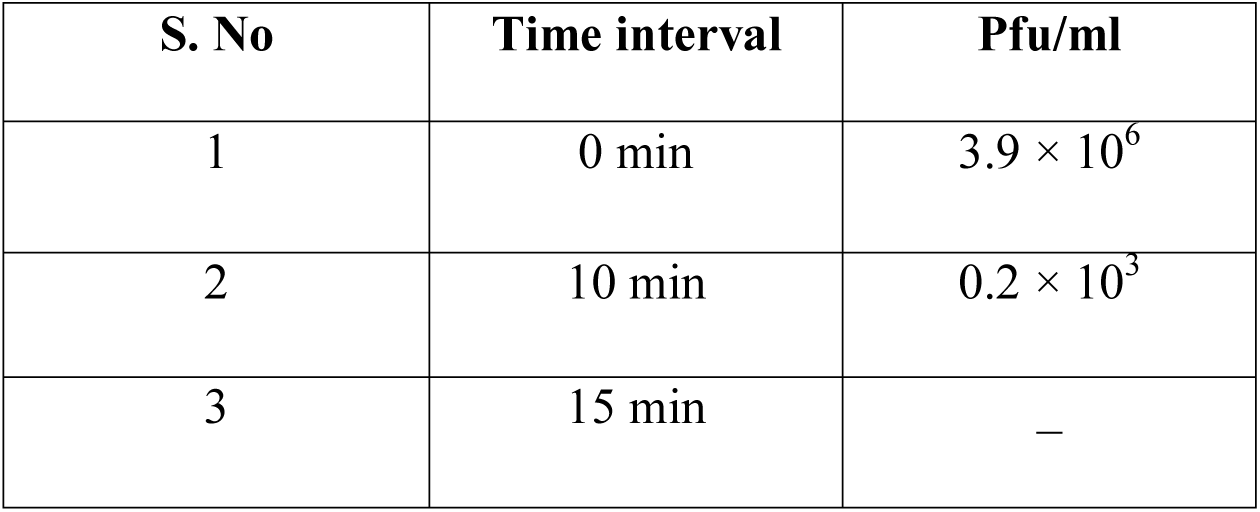
Effect of chloroform on phage survival.

**Fig 5:**
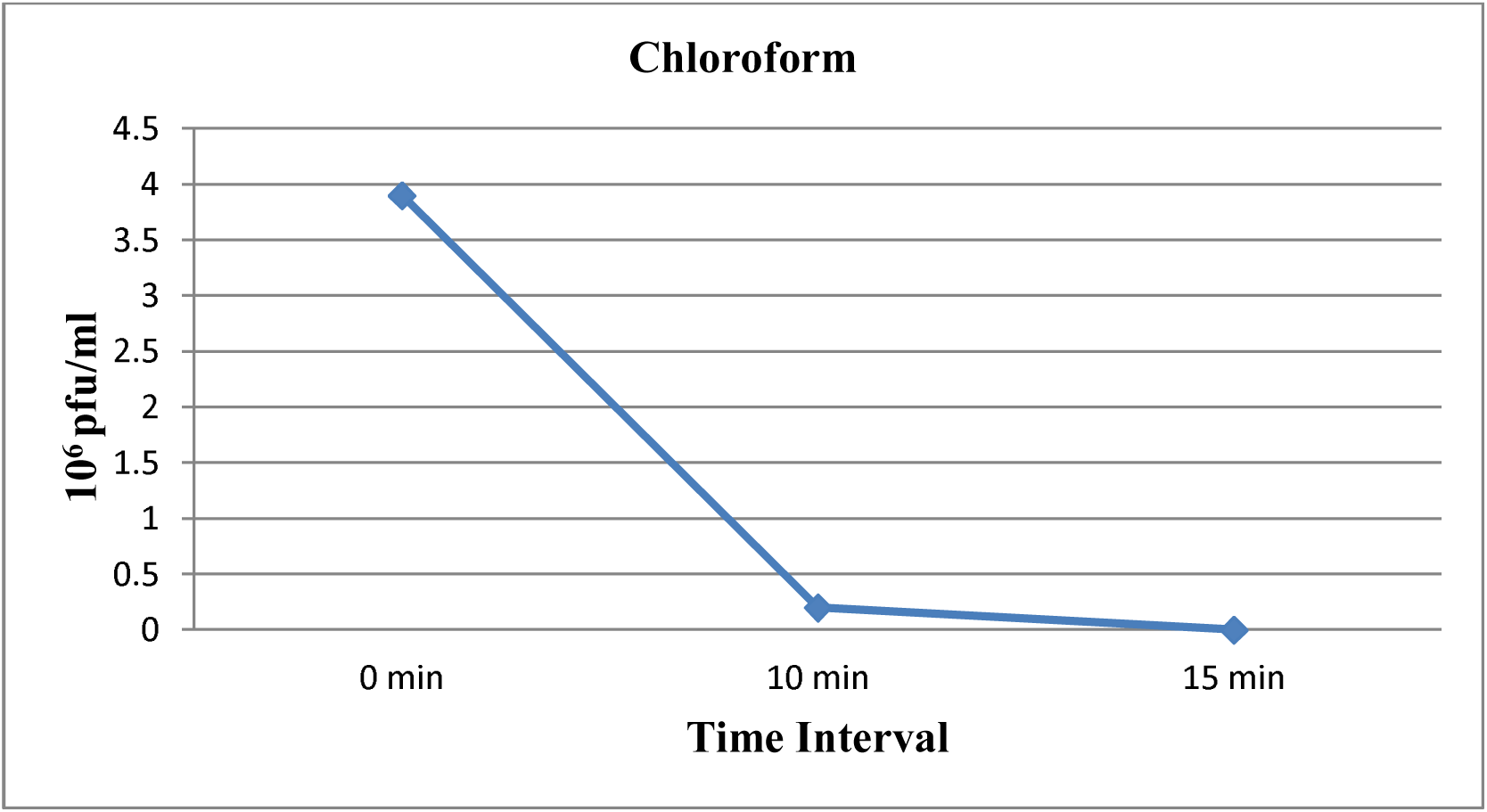
Effect of chloroform on phage survival.

### Effect of SDS on phage

The effect of sodium dodecyl sulphate (SDS) treatment on activity of phage was studied. It was found that both 1% and 0.1% concentrations of SDS completely inactivated the phage within 15 minutes at 37°C (Table 6, fig6). Pandey *et al* (2013) had also reported complete inactivation of phage within 15 minutes of SDS treatment.

**Table 6:** Effect of SDS treatment on phage.

| S. No | Time interval | Pfu/ml in<br>0.1% SDS | Pfu/ml in<br>1.0% SDS |
| --- | --- | --- | --- |
| 1 | 0 min | $4.4 \times 10^6$ | $4.2 \times 10^6$ |
| 2 | 10 min | $0.3 \times 10^6$ | $0.1 \times 10^6$ |
| 3 | 15 min | – | – |

**Fig 6:**
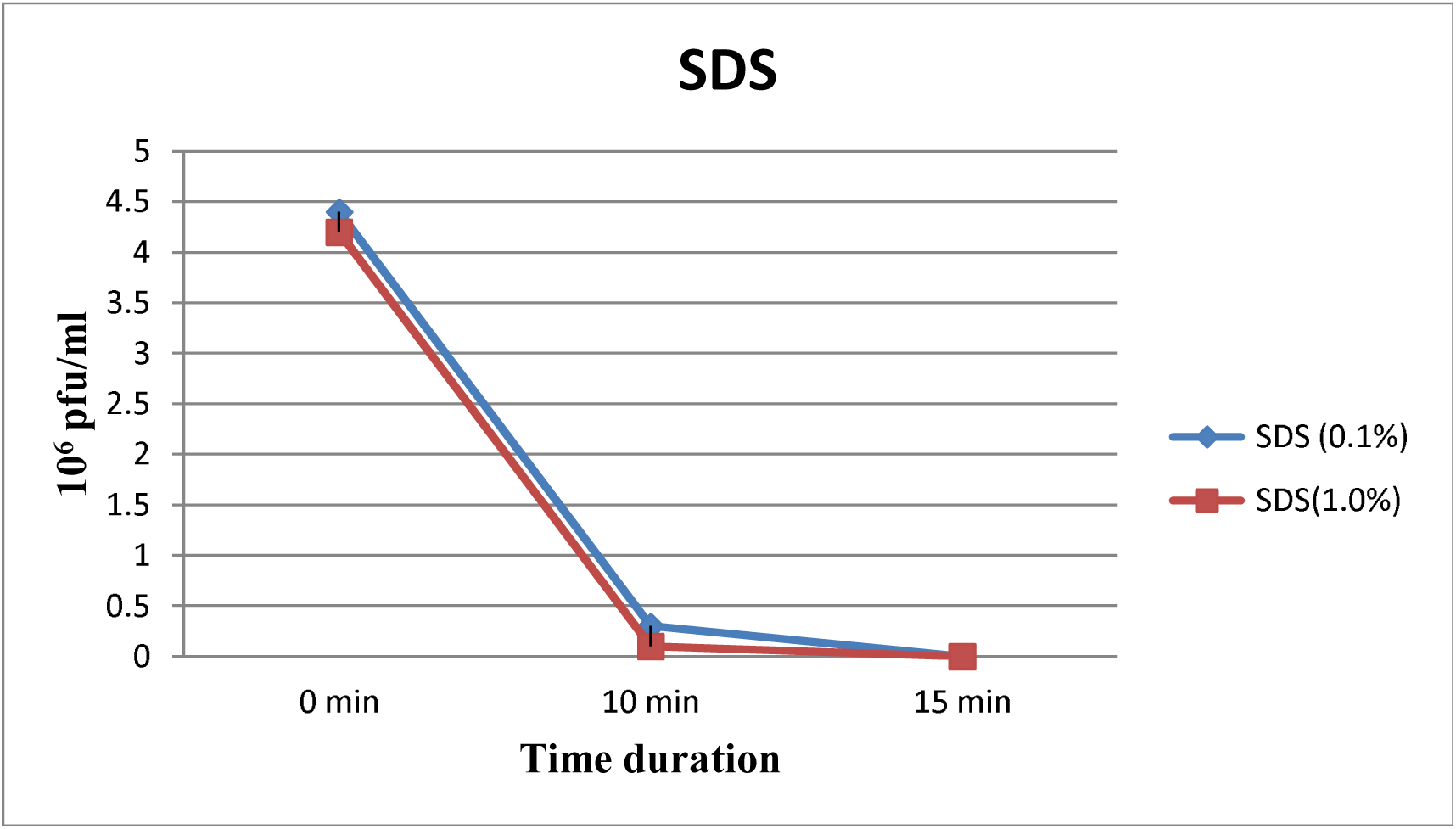
Effect of SDS on phage.

## Effect of enzymes on phage activity

### Effect of lysozyme

Lysozyme at a concentration of 20 mg/ml completely inactivated the phage within one hour (Table 7, fig 7). Pandey *et al* (2013) had also reported complete inactivation of their phage within 1 hour with lysozyme treatment.

**Table 7:** Effect of lysozyme on phage.

| S. No | Duration of exposure | Phage count (Pfu/ml) |
| --- | --- | --- |
| 1 | 0 min | $3.5 \times 10^6$ |
| 2 | 10 min | $1.0 \times 10^6$ |
| 3 | 30 min | $0.1 \times 10^6$ |
| 4 | 60 min | — |

**Fig 7:**
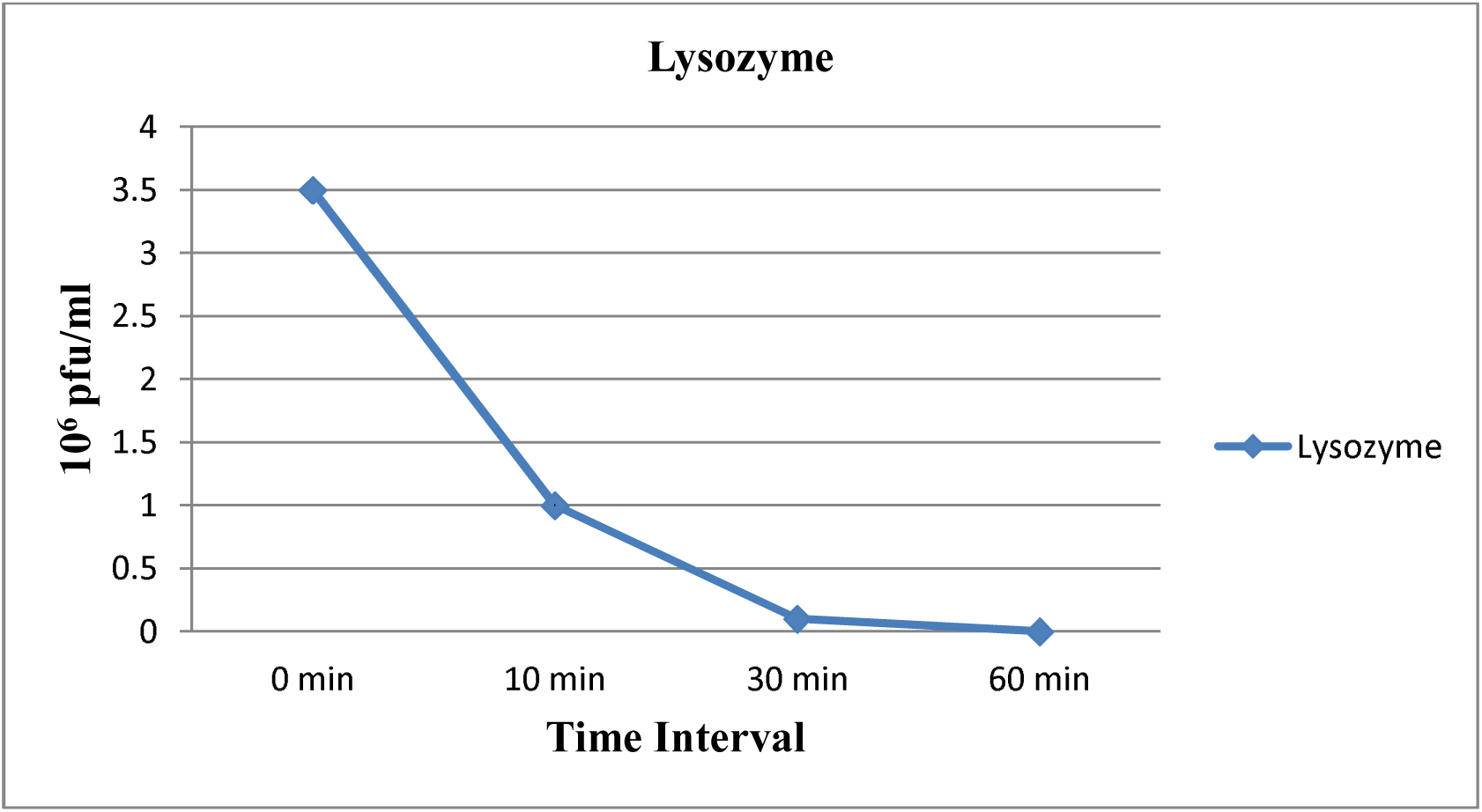
Effect of lysozyme on phage.

### Effect of RNase on phage

No detectable change was found on treatment of phage with RNase for 120 minutes (Table 8, fig 8). Morris *et al* (1973) had also reported no detectable effect on phage titre after RNase treatment for 90 min. Pandey *et al* (2013) had reported that phage remains almost stable up to 3 hours after treatment with RNase.

**Table 8:** Effect of RNase on phage.

| S. No | Duration of exposure | Phage count (Pfu/ml) |
| --- | --- | --- |
| 1 | 0 min | $4.4 \times 10^6$ |
| 2 | 10 min | $4.2 \times 10^6$ |
| 3 | 30 min | $3.9 \times 10^6$ |
| 4 | 60 min | $3.5 \times 10^6$ |
| 5 | 120 min | $2.5 \times 10^6$ |

**Fig 8:**
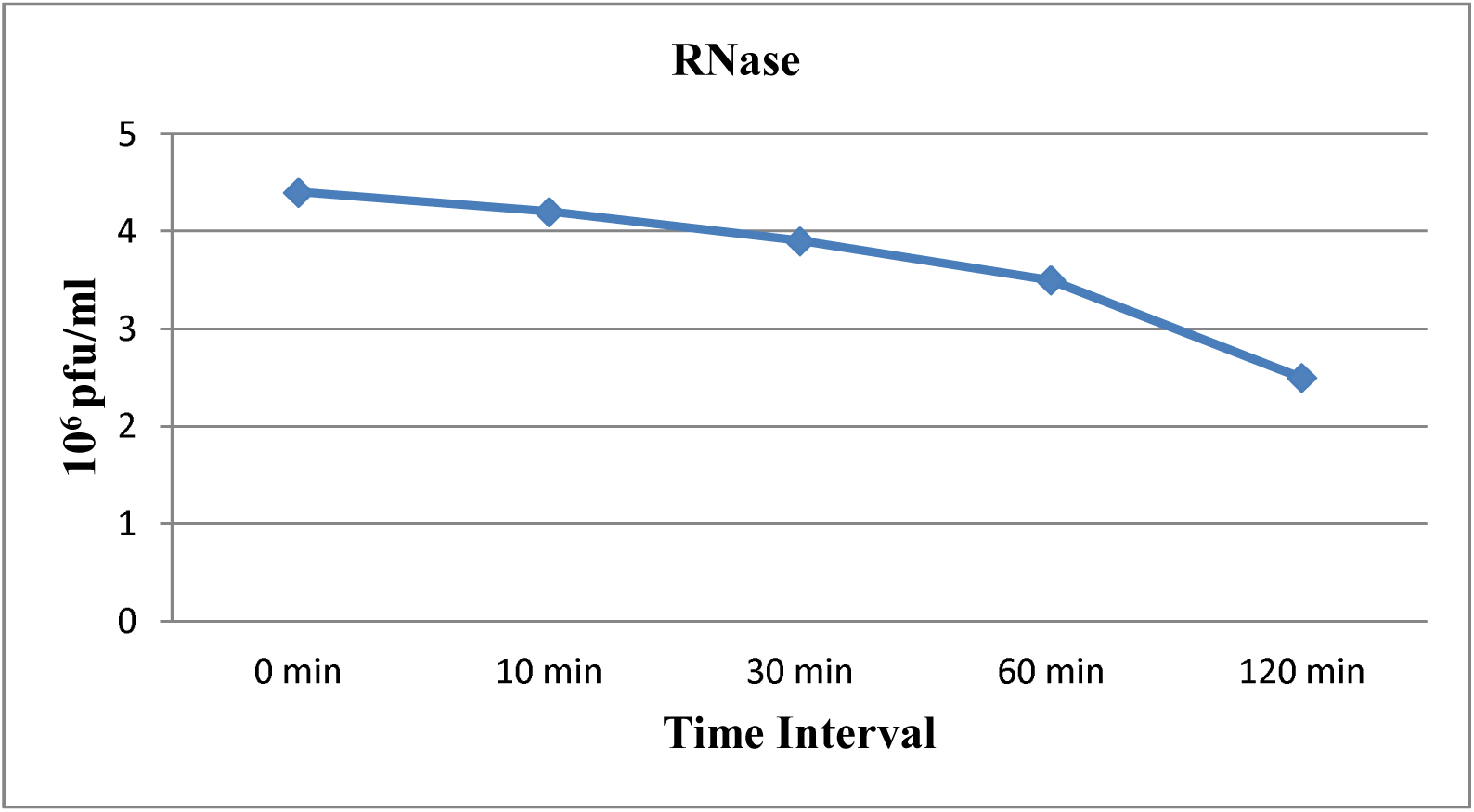
Effect of RNase on phage.

**Fig 4:**
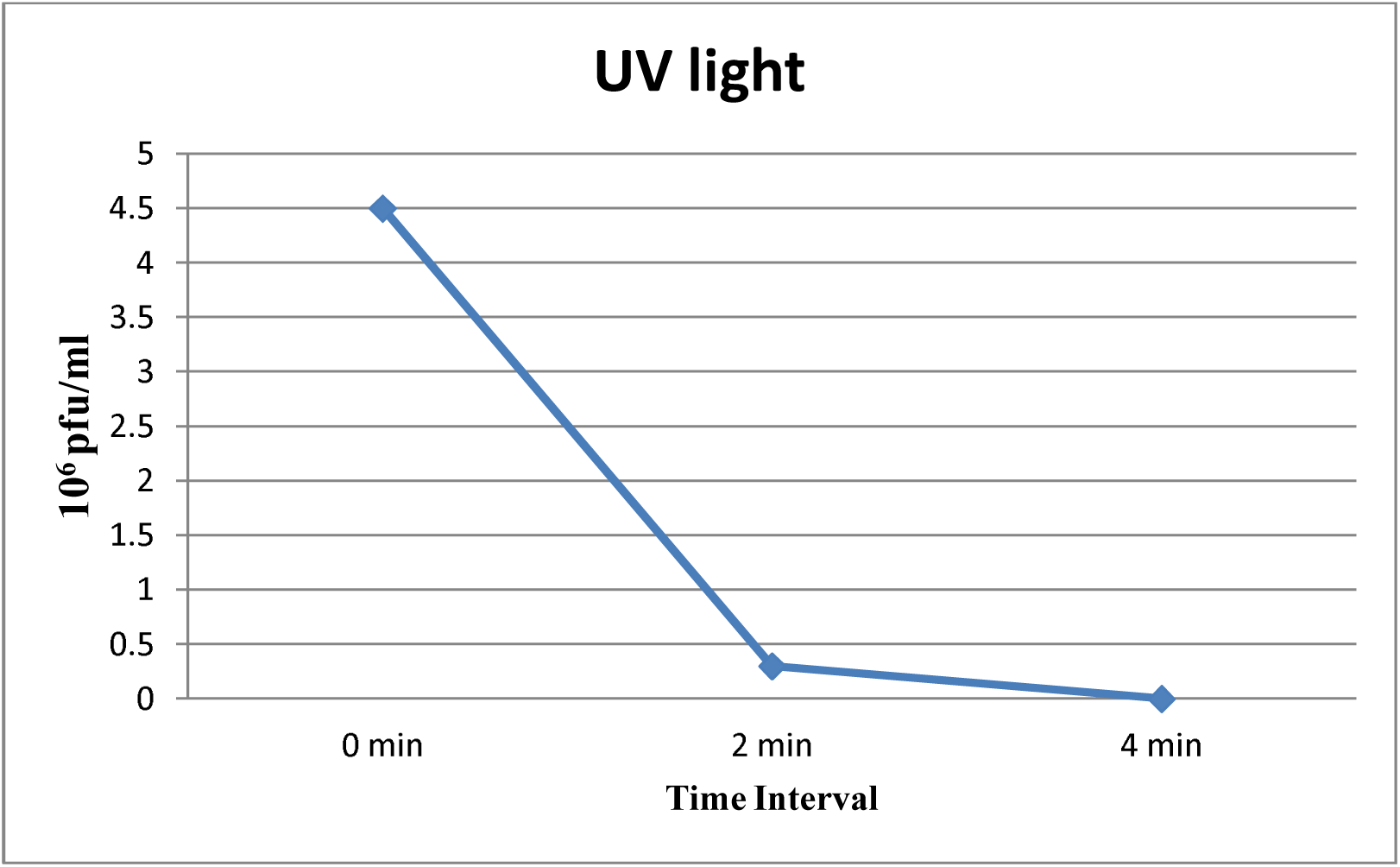
Effect of exposure to UV light on phage survival.

**Table 4:** Effect of UV light on brucellaphage.

| S. No | Time duration of exposure | Phage count (Pfu/ml) |
| --- | --- | --- |
| 1 | 0 min | $4.5 \times 10^6$ |
| 2 | 2 min | $0.3 \times 10^6$ |
| 3 | 4 min | – |

